# ITS-based assessment of Madagascar’s fungal diversity and arrival of ectomycorrhizal fungi to the island

**DOI:** 10.1101/2022.03.09.483579

**Authors:** Mauro Rivas-Ferreiro, Laura M. Suz, Shannon M. Skarha, Franck Rakotonasolo, Bryn T.M. Dentinger

## Abstract

Madagascar is known for its high diversity and endemism of Fauna and Flora, which makes it particularly interesting for research on diversity and evolution. Fungi, however, have been largely overlooked, and whether fungi exhibit the same patterns as animals and plants has yet to be further examined.

We collected fungal sporocarps and ectomycorrhizal (EcM) roots during opportunistic surveys in five forests in Madagascar and generated a dataset of fungal Internal Transcribed Spacer (ITS) DNA sequences. We analysed them together with all publicly available fungal ITS DNA sequences. We identified 620 Operational Taxonomic Units (OTUs) from Madagascar, 10% of which contained only sequences from our surveys. We found 292 OTUs belonging to EcM species with /russula-lactarius, /boletus, /tomentella-telephora, /cortinarius and /amanita as the most abundant EcM lineages. Overall, sixty percent of all the fungi and 81% of the EcM species found are endemic to Madagascar. Lastly, we conducted a phylogenetic analysis using all the OTUs in Amanitaceae, Boletaceae and Russulaceae families to elucidate their relative timing of arrival in Madagascar. We found that the EcM species from Madagascar in the three families diverged recently (less than 34 mya), long after the separation of India and Madagascar (88 mya), which is consistent with a dispersal mediated process of arrival on the island. Our study provides the first comprehensive view of the current state of knowledge of EcM fungi in Madagascar represented by molecular data useful for ecological and evolutionary studies.

## Introduction

According to the Convention on Biological Diversity (CBD 2014), Madagascar is a megadiverse country with a high number of endemic species. This is, however, based entirely on plant and animal studies, since there is very little literature regarding its Funga (Buyck 2008), and as a result, not much is known about the current fungal diversity in Madagascar (Ralaiveloarisoa *et al*. 2020).

The first scientific reports based on sporocarp collections for the Funga of Madagascar, date back to the late 19^th^ century (Cooke and Massee 1890). After several studies conducted by French mycologists in the first half of the 20^th^ century, including those of Patouillard (1924, 1928), Romagnesi (1941), Métrod (1949) and Heim (1936, 1938a, b, 1945), there was a large gap from World War II onwards when the fungal diversity of the island was not studied (Vizzini *et al*. 2019). It was not until very recently that the interest in Madagascar fungi resurfaced, with several new species being reported and monographs being published (Eyssartier and Buyck 1999 a, b; Eyssartier *et al*. 2001; Antonín *et al*. 2005; Antonín and Buyck 2006; Duhem and Buyck 2007; Buyck *et al*. 2007; Buyck, 2008; Aptroot 2016; Shay *et al*. 2017; Ralaiveloarisoa *et al*. 2020, 2021).

According to the latest National Report of the Convention on Biological Diversity, a total of 292 fungal species are currently inventoried for Madagascar (Madagascar Ministry of Environment and Forests 2014). Some authors report about 500 sporocarp-forming fungi collected from its forests, with 90% of them still waiting to be described as they could be new to science (Ralaiveloarisoa *et al*. 2020). In a recent study, Malagasy mycologists collected more than 1,600 sporocarps from the Analamazaotra Special Reserve (Andasibe, Madagascar) that were classified in 165 morphospecies (defined by grouping sporocarps based on their distinct morphological characters) from which only 19 could be identified at species level, indicating that more than 85% of the morphospecies collected in this Reserve could belong to new, undescribed taxa (Ralaiveloarisoa *et al*. 2016).

The island of Madagascar contains an immense variety of ecosystems, ranging from the dry deciduous forests of the West, the Central highlands with some areas of ericoid thicket throughout, to the evergreen rainforests of the East, with spiny thickets in the southern part of the island and Sambirano rainforest in the North (Yoder and Nowak 2006). Because most Malagasy organisms evolved in isolation, numerous evolutionary radiations with microendemisms emerged as many species adapted to these pronounced environmental gradients (Vences *et al*. 2009). Plants, with 83% of the species being endemic, are one of such highly diverse and highly endemic groups of organisms (Goodman and Benstead 2005). Notably, some of the endemic families in Madagascar, such as Sarcolaenaceae and Asteropeiaceae, form obligate ectomycorrhizal (EcM) relationships with fungi (Ducousso *et al*. 2008).

Ectomycorrhizal fungi (mostly belonging to the Basidiomycota phylum, but also to the Ascomycota and Mucoromycota phyla) are one of the most prominent ecological guilds in forest ecosystems. They form symbiotic relationships with the roots of about 30 plant lineages worldwide (Brundrett 2002, Ducousso *et al*. 2008, Tedersoo *et al*. 2020), in which the fungus obtains plant sugars from photosynthesis in exchange for mineral nutrients and water gathered from the soil (Smith & Read, 2008). EcM fungal hyphae that ensheath the root tips is also thought to offer protection to the host plants (Tedersoo *et al*. 2020). The EcM lifestyle has evolved at least 80 times in different fungal lineages, a testament to its success as a lifestyle for fungi (Brundrett 2002, Tedersoo and Smith 2013).

Although research on the EcM symbiosis in tropical ecosystems has increased over the past 50 years (Ducousso *et al*. 2008, 2012; Henry *et al*. 2017), the diversity of tropical EcM fungi is still poorly known (Brundrett 2002, Tedersoo and Smith 2013). In particular, the mycorrhizal communities associated with native forests in Madagascar have not been thoroughly studied (Henry *et al*. 2015). In Madagascar, the EcM lifestyle is not restricted to endemic plant families; other non-endemic tropical plant species such as *Intsia bijuga* (Fabaceae) and several species of *Uapaca* (Phyllanthaceae), as well as exotic trees of the genera *Pinus* and *Eucalyptus*, are also EcM (Buyck 2002, Ducousso *et al*. 2008, 2012).

How and when the EcM lifestyle arrived in Madagascar has been unresolved for some time (Buyck 2002). Madagascar has a very particular geological history: the island was once a part of the supercontinent Gondwana, forming a land mass with India that started drifting from the supercontinent about 100 mya, until their final separation 88 mya (Yoder and Nowak 2006). This has led to many hypotheses about the speciation events that might have led to its high rates of endemism across kingdoms, and how certain lifestyles such as the EcM fungi emerged in the island. Theories about the arrival of the EcM symbioses in Madagascar are largely based on the evolutionary history of their plant partner, with two main hypotheses proposed so far.

The first hypothesis, based in Ducousso *et al*. (2004), suggests a vicariance-driven origin of the EcM habit in Madagascar based on phylogenetic analyses showing that the endemic EcM family Sarcolaenaceae shared a most recent common ancestor with the EcM Dipterocarpaceae, with representatives in Asia, Africa and South America. A parsimonious reconstruction of the EcM condition in these two families places a single gain in their common ancestor, somewhere between 165-155 mya (Yoder and Nowak, 2006), prior to the disarticulation of Gondwanaland (Ducousso et al. 2004). However, in addition to the dubious assumptions about the evolutionary likelihood of independent origins of the EcM habit, important gaps in the knowledge of the phylogenetic relationships between Sarcolaenaceae and Dipterocarpaceae, as well as among the dipterocarp clades, themselves (APG 2016), question this hypothesis.

The second hypothesis states that the arrival of the EcM habit occurred more recently, with the dispersal of the EcM trees from continental Africa and other areas. Several authors have suggested through phylogenetic analyses that many EcM plant species that occur in Madagascar arose quite recently (Zanne *et al*. 2014, Janssens *et al*. 2020); for some of them, long distance dispersal is also known to occur (Lewis *et al*. 2005).

As seen in other biogeographic studies, both hypotheses are not mutually exclusive (Peterson *et al*. 2010, Sauquet *et al*. 2012); due to the polyphyletic nature of the EcM symbiosis in the fungal tree of life, the origin of this symbiosis in Madagascar could be explained by either or both hypotheses, and it could also be different in different clades. To fully understand the diversity and origin of EcM fungi in Madagascar, a comprehensive baseline of the fungal diversity in the island is needed from which accurate and reliable inferences can be drawn.

In an effort to establish a diversity baseline for fungi in Madagascar, we had the following aims: 1) to explore the potential of opportunistic field surveys to document Malagasy fungi, 2) to compile the known Malagasy fungal diversity, using all available fungal ITS DNA sequence data and a clustering approach to define species, 3) to explore patterns of EcM diversity in Malagasy fungi, and 4) to investigate the phylogenetic patterns in three dominant EcM fungal families to elucidate the mechanism of arrival of the EcM lifestyle in Madagascar.

## Materials and methods

### Opportunistic field surveys and new fungal ITS sequences generated in this study

Opportunistic sampling of sporocarps of macrofungi and root samples from six EcM host genera (the Madagascar endemic *Sarcolaena, Leptolaena* and *Asteropeia*, the African-Malagasy *Uapaca*, the Indo-Pacific *Intsia* and the Pacific *Eucalyptus*) was carried out in five sites in Madagascar during three field trips in 2012, 2014 and 2017 (Table 1). Habitats included eastern humid forest (Saha Forest Camp, Vohimana, Antsinanana), littoral forest (Mandena) and dry western forest (Ankarafantsika).

**Table 1.**
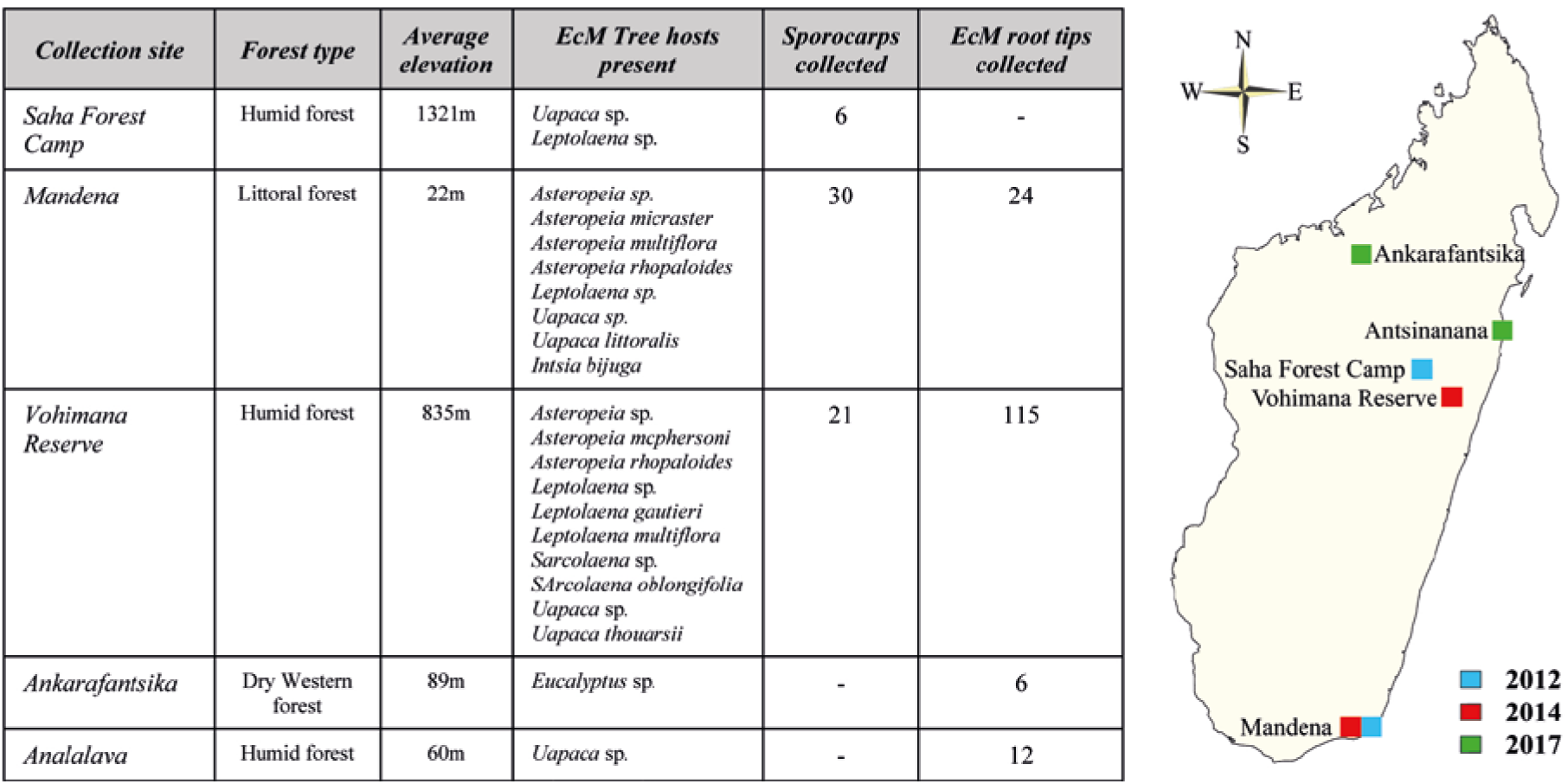
Study sites in Madagascar where the collection of sporocarps and ectomycorrhizal roots were conducted, and the map showing the geographical and chronological distribution of the sampling campaigns.

When present, sporocarps were photographed and collected (Fig. 1) and root samples from the nearest EcM tree were collected underneath. Sporocarps were dried in silica gel or in a food drier for preservation. Mycorrhizal root samples were collected from 67 EcM trees (Table 1) by tracking lateral roots from the main trunk and stored in plastic bags. Roots were rinsed with water in a 0.5 mm soil sieve on the same day of collection and stored in 2X CTAB buffer until further processing.

**Fig. 1.**
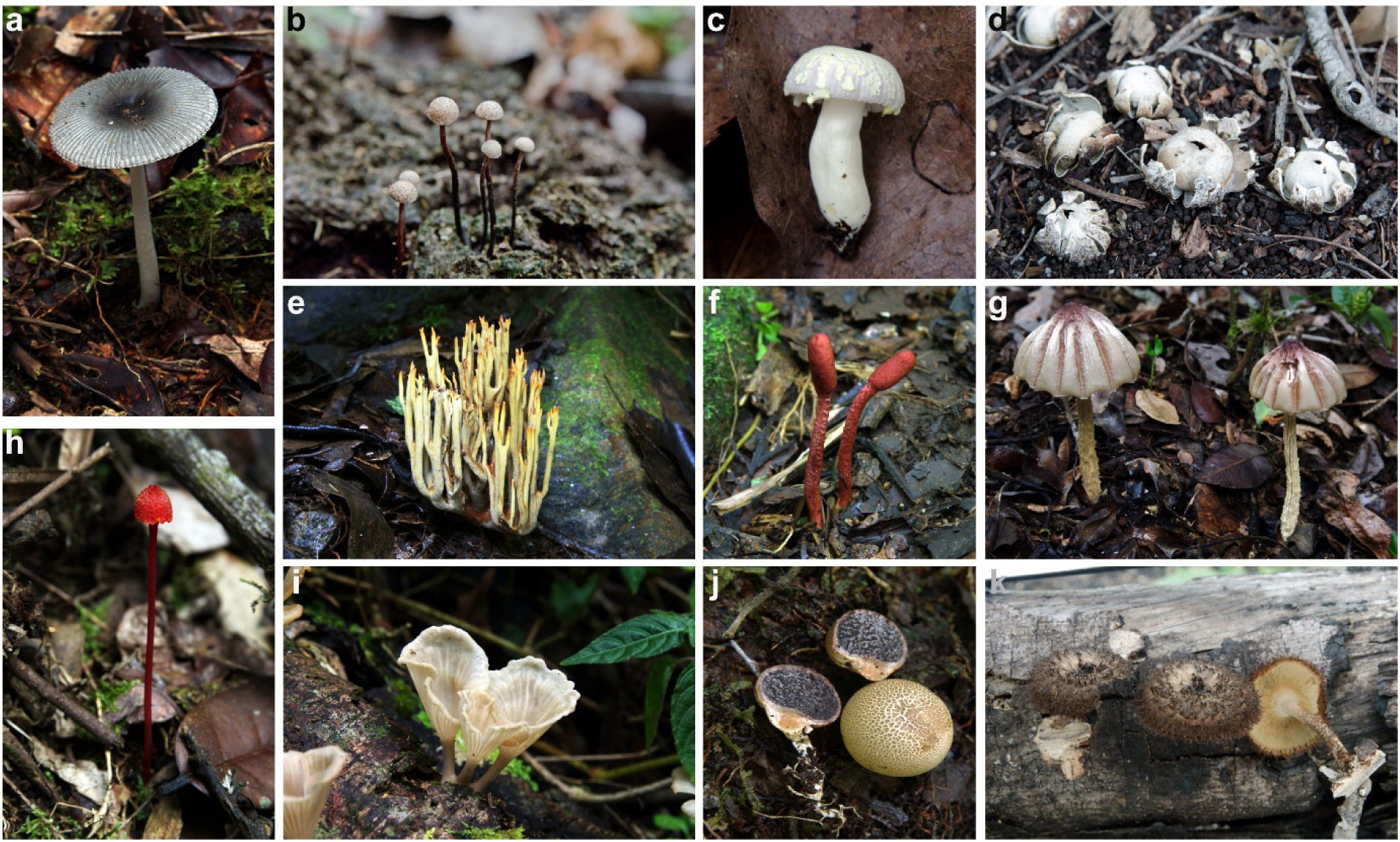
Sporocarps collected in 2012 in Mandena (b, c, d, k) and in 2014 in Vohimana Reserve (a, e, f, g, h, i, j); a- *Amanita* sp., b- *Poronia* sp., c- *Russula* sp., d- *Astraeus* sp., e- *Ramaria* sp., f- *Cordyceps* sp., g- *Marasmius* sp., h- *Hygrocybe* sp., i- *Trogia* sp., j- *Scleroderma* sp., k- *Lentinus* sp.

Once in the laboratory, roots were observed under a dissecting microscope for the presence of EcM root tips. When possible, eight EcM roots were selected and one ectomycorrhiza from each root was randomly collected for DNA analysis.

Genomic DNA was isolated using ∼1 mg of hymenophore tissue in 20 μL of Extract-N-Amp (Sigma) for sporocarps, and 10 μL for the individual ectomycorrhizal root tips. The forward ITS1F (Gardes and Bruns 1993) and reverse ITS4 (White *et al*. 1990) primers were used to amplify the internal transcribed spacer (ITS) region of the nrDNA, through PCR. PCR products were purified using ExoSAP-IT (USB, Cleveland, OH, USA) and sequenced bidirectionally using BigDye3.1 in an ABI 3730 (Applied Biosystems, Foster City, CA, USA). The obtained sequences were manually edited using Sequencher v. 4.2. (Gene Codes Corp., Ann Arbor, MI, USA).

### Fungal ITS sequences from PlutoF, GenBank and UNITE

To retrieve all the fungal ITS sequences publicly available, we used the *Full UNITE+INSDC* fungal dataset (version 8.3, with 1039010 sequences) available in UNITE (Abarenkov *et al*. 2010a). This dataset contains all curated ITS sequences in the UNITE and INSDC databases, that belong to a UNITE Species Hypotheses (Abarenkov *et al*. 2021a).

To retrieve all complete or partial ITS sequences from fungi collected in Madagascar, we carried out a search in *PlutoF* (Abarenkov *et al*. 2010b, last accessed on 07/June/2021) and *GenBank Nucleotide* databases (NCBI 1988, last accessed on 07/June/2021) For PlutoF, we used the filters “Taxon: Fungi R.T. Moore, 1980”, “Country: Madagascar” and “Include subtaxa: Yes”; in GenBank, we used the search “country=[All Fields] AND “Madagascar”[All Fields] AND fungi[filter]”

### Clustering analysis

To standardise sequence lengths and remove the unwanted LSU and SSU regions, we processed the ITS sequences using the software *ITSx* (Bengtsson-Palme *et al*. 2013). To remove all duplicate sequences, the datasets containing PlutoF and GenBank sequences from Madagascar were combined and dereplicated using the *--derep_fulllength* option of the vsearch software (Rognes *et al*. 2016). The resulting dataset (PlutoF+GenBank dataset) was then used as a representation of the known fungal ITS diversity of Madagascar.

Taxonomic assignment of the newly generated sequences and the PlutoF+GenBank dataset from Madagascar was performed using the *assignTaxonomy* function from the R package *dada2* (Callahan *et al*. 2016), using the *UNITE General FASTA release* (version 8.3, 98.084 sequences) as a reference database (Abarenkov *et al*. 2021b). The resulting annotated sequences from Madagascar were then combined with the UNITE dataset (all fungi) and the clustering analysis was carried out using *vsearch* (Rognes *et al*., 2016) with a 97% similarity threshold to delimit Operational Taxonomic Units (OTUs). These OTUs were used as a proxy for species in further analysis; when referring to the OTUs as biological entities, we will use the term “species”. For the graphical expression of the OTU’s taxonomic diversity, we used the R package *metacoder* (Foster et al. 2017).

To assign the EcM status and EcM lineage to the OTUs that contained sequences with a Species Hypothesis (SH) number (Kõljalg et al. 2013), we used the information available in the UNITE database. For those OTUs that did not contain sequences with SH numbers, we used *FUNGuild* (Nguyen *et al*. 2016), which assigned a guild to each OTU based on the taxonomy provided by *dada2*, and confirmed which species were EcM by consulting the available literature (Tedersoo *et al*. 2010, Tedersoo and Smith 2013).

### Phylogenetic analysis

We generated three family-level datasets with the OTUs obtained in our clustering analyses belonging to the EcM families Amanitaceae, Boletaceae and Russulaceae, as they are among the most abundant EcM families found in tropical forests (Ramanankierana *et al*. 2007; Henkel *et al*. 2012; Corrales *et al*. 2018), to reconstruct their phylogenetic histories and evaluate the two main hypotheses for the arrival of the EcM fungi to the island.

Since the resulting family datasets were quite large, to reduce computational efforts without losing the information about the Malagasy species, we de-replicated and reclustered only the UNITE sequences from each family using the algorithm *h-dh-hit-est* (three runs, with a sequence identity cut-off value of 0.99, 0.98 and 0.97, respectively), available in the *CD-HIT Suite* (Huang *et al*. 2010). The resulting OTUs were then manually explored, and dubious OTUs (potential misidentifications or OTUs with many sequences from other families) were removed from the analysis. This resulted in three datasets (one for each EcM family) with one representative UNITE sequence per OTU. These revised datasets were then merged again with the Malagasy sequences, which were not part of this second clustering process in order to preserve all the available information about the Malagasy species.

The new family-level datasets were aligned using the *FFT-NS-2* algorithm in *MAFFT v7*.*475* (Katoh 2013), and the ITS1, 5.8S and ITS2 regions were manually identified. Then, a maximum likelihood analyses was carried out in *IQ-TREE* (Nguyen *et al*. 2015) using a partitioned analysis, with the manually identified ITS1, 5.8S and ITS2 regions as partitions (Chernomor *et al*. 2016), applying *ModelFinder* to find the best substitution model for each partition (Kalyaanamoorthy *et al*. 2017) and the *Ultrafast Bootstrap* approximation (Hoang *et al*. 2018) with 1000 replicates.

After the maximum likelihood trees were obtained for each EcM family, an ultrametric tree was generated using a penalised likelihood approach with uncorrelated branch lengths, using the *chronos* function from the R package *ape* version 5.5 (Paradis 2013; Paradis *et al*. 2019) in R Studio Version 1.4.1717 (R Core Team 2021). Due to the lack of EcM fungal fossils, we used a relative dating approach using the mean stem age for Amanitaceae, Boletaceae and Russulaceae described in He *et al*. (2019) as the family divergence times (125 mya, 98 mya and 97 mya, respectively).

## Results

### Sporocarp and EcM root tip DNA sequencing

We generated 214 sequences, 58 corresponding to sporocarps and 156 to ectomycorrhizae. All sequences are available in GenBank under the accession numbers OM219812-OM219816 and OM219818-OM220026; additional information on the sequences can be found in Table S1.

### Fungal DNA sequences in public databases

We recovered 2,467 ITS sequences from fungi collected in Madagascar from GenBank and 1,652 sequences from the PlutoF database. After combining the datasets, extracting the ITS region and removing duplicates, the final dataset (PlutoF+GenBank) was made up of 1,179 publicly available sequences from Madagascar. From the UNITE dataset, after ITS extraction and dereplication, we obtained a total of 735,598 ITS sequences out of the original 1,039,010.

### Clustering analysis

#### Fungal diversity of Madagascar

We recovered a total of 620 fungal OTUs present in Madagascar (Fig. 2; Table S2). From these, 545 contained previously published ITS sequences from Madagascar; the remaining 75 OTUs were novelties to the fungal diversity of the island. Basidiomycota is the most abundant phylum, containing 402 OTUs with Russulaceae (87 OTUs, 67 of which correspond to the genus *Russula*), Thelephoraceae (41 OTUs) and Boletaceae (31 OTUs), followed by Cortinariaceae (22 OTUs) and Amanitaceae (15 OTUs) as the most abundant families.

**Fig. 2.**
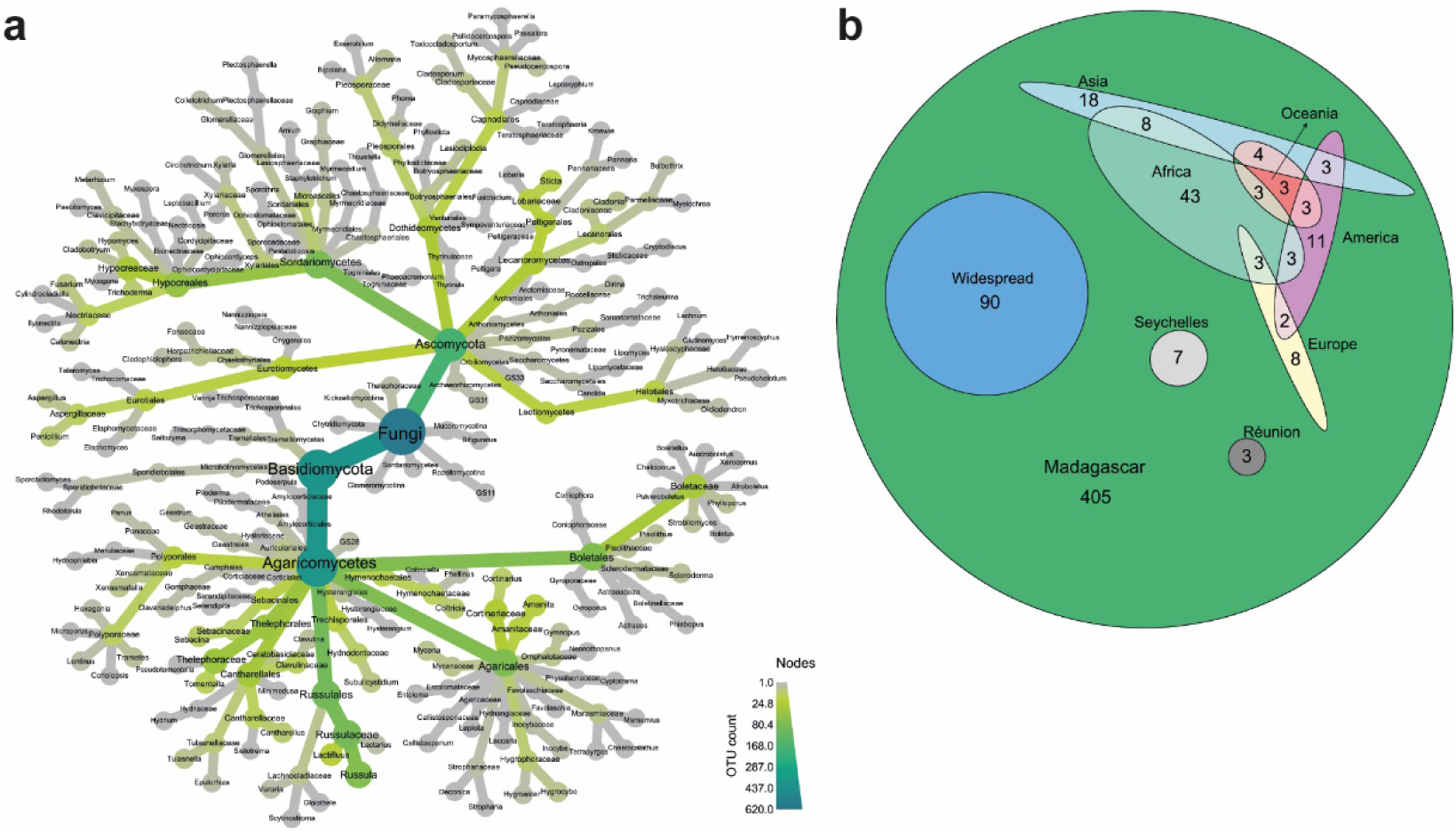
a- Metacoder chart showing the taxonomic diversity of the 620 Operational Taxonomic Units (OTUs) obtained in the clustering analysis of the Internal Transcribed Spacer (ITS) sequences from Madagascar; b- Euler diagram including the number of OTUs found in Madagascar and their global distribution.

One hundred and seventy-seven species belonged to the phylum Ascomycota with the families Hypocreaceae (29 OTUs), Lobariaceae (14 OTUs), Nectriaceae (10 OTUs) and Aspergillaceae (10 OTUs) being the most abundant. We also found Kickxellomycotina (two OTUs), Rozellomycotina, Chytridiomycota, Mucoromycotina and Glomeromycotina (one OTU each) while 35 OTUs could not be identified to phylum level (Fig. 2A). Only 138 OTUs (22%) out of the 620 contained a sequence identified at species-level (either from a UNITE Species Hypothesis, or from the assignment by *dada2*).

Two thirds of the fungal diversity from Madagascar found in our clustering analysis (405 OTUs, 65% of the total) contained sequences referenced only for Madagascar and therefore were considered endemic. Some species were referenced for Madagascar and the islands of Seychelles (seven OTUs) and Réunion (three OTUs). We also found species present only in Madagascar and continental Africa (43 OTUs), species present in Madagascar, Africa and one other continent (17 OTUs), while some others were present in Madagascar and another continent, but not continental Africa (55 OTUs) (Fig. 2B).

The remaining 90 OTUs were found in three or more continents, considered in this study as widespread (Fig. 2B). More than two thirds of the widespread OTUs belong to Ascomycota, mainly to Sordariomycetes (29 OTUs), Dothideomycetes (20 OTUs) and Eurotiomycetes (11 OTUs).

Of the 75 species that were not previously found in Madagascar, 14 were previously found elsewhere: five of those OTUs contained widespread species, while three OTUs were only known from the Americas, two OTUs only from Asia, two OTUs from Africa and Asia, one OTU from America and Oceania, and one OTU from Africa and Oceania. The remaining 61 species contained only sequences newly generated from our opportunistic sampling campaigns in Madagascar. These species, potentially new to Madagascar, contained mostly sequences from ectomycorrhizae (60%; 45 newly described OTUs), while the remaining contained sequences from sporocarps (36%; 27 OTUs) or from both EcM roots and sporocarps (4%; three OTUs).

### EcM fungal diversity of Madagascar

Two-hundred ninety-two OTUs (47% of the total fungal species of Madagascar) found in our clustering analysis were classified as EcM. Among these species, the main EcM lineages were /russula-lactarius, /boletus, /tomentella-telephora, /cortinarius and /amanita (Fig. 3A). Of those, 236 OTUs (81% of the EcM species found) were endemic, containing sequences referenced only for Madagascar. Thirty-six of the remaining OTUs were found in Madagascar and continental Africa, 6 in Madagascar and the Seychelles and 1 in Madagascar and Réunion, as well as Madagascar, Africa and Oceania (2 OTUs), Madagascar, America and Oceania (1 OTU), Madagascar and Oceania (3 OTUs), and Madagascar, Africa and Asia (1 OTU). Six OTUs were widespread (Fig. 3B).

**Fig. 3.**
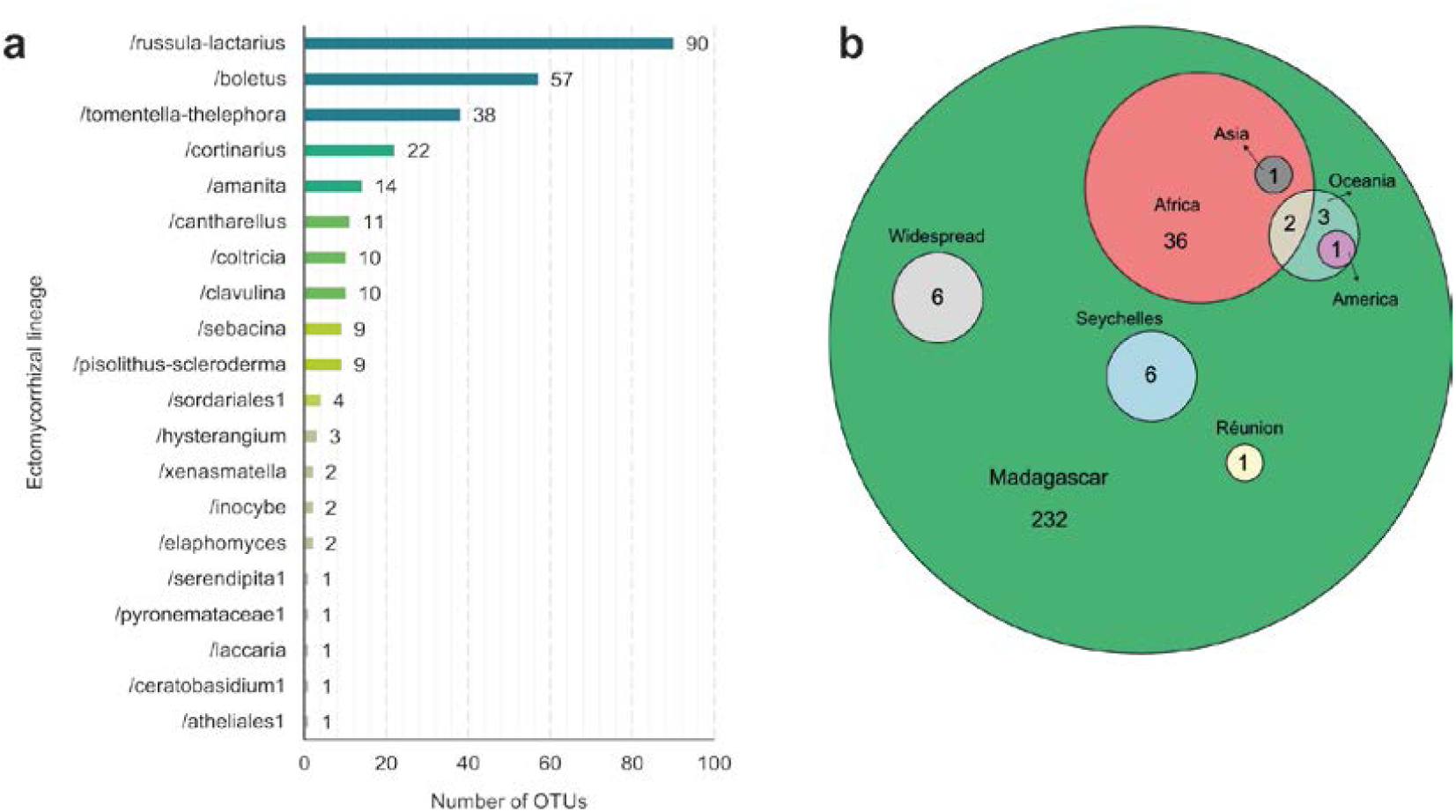
a- Diversity of ectomycorrhizal (EcM) lineages of the 292 Operational Taxonomic Units (OTUs) found from Madagascar; b- Euler diagram including the number of EcM OTUs found in Madagascar and their global distribution.

Of the total EcM diversity found, 20 OTUs contained only new sequences generated in this study. Of those, 12 OTUs contained exclusively sequences from EcM roots, 6 contained sequences from sporocarps, and 2 contained both sporocarp and EcM root sequences. They belonged to the EcM lineages /russula-lactarius (9 OTUs), /boletus (5 OTUs), /amanita (2 OTUs), /cantharellus, /elaphomyces, /sebacina and /cortinarius (1 OTU each). These newly reported species account for 7% of the current known EcM fungal diversity in Madagascar.

### Phylogenetic analyses

From the clustering analysis, we extracted 984 OTUs (5,485 sequences), 1,107 OTUs (5,754 sequences) and 2,925 OTUs (5,485 sequences) corresponding to the families Amanitaceae, Boletaceae and Russulaceae, respectively. After re-clustering, selecting one representative sequence per OTU, deleting dubious OTUs, and merging with the Madagascar sequences, the Amanitaceae, Boletaceae and Russulaceae datasets were made up of 952, 1,232 and 3,083 sequences, respectively.

#### Amanitaceae

Five of the Amanitaceae sequences from Madagascar used in the phylogenetic reconstruction were generated in this study, while 19 were obtained from public repositories. Our relative dating of the Amanitaceae family (Fig. 4A) showed that all species from Madagascar diverged less than 34 mya.

**Fig. 4.**
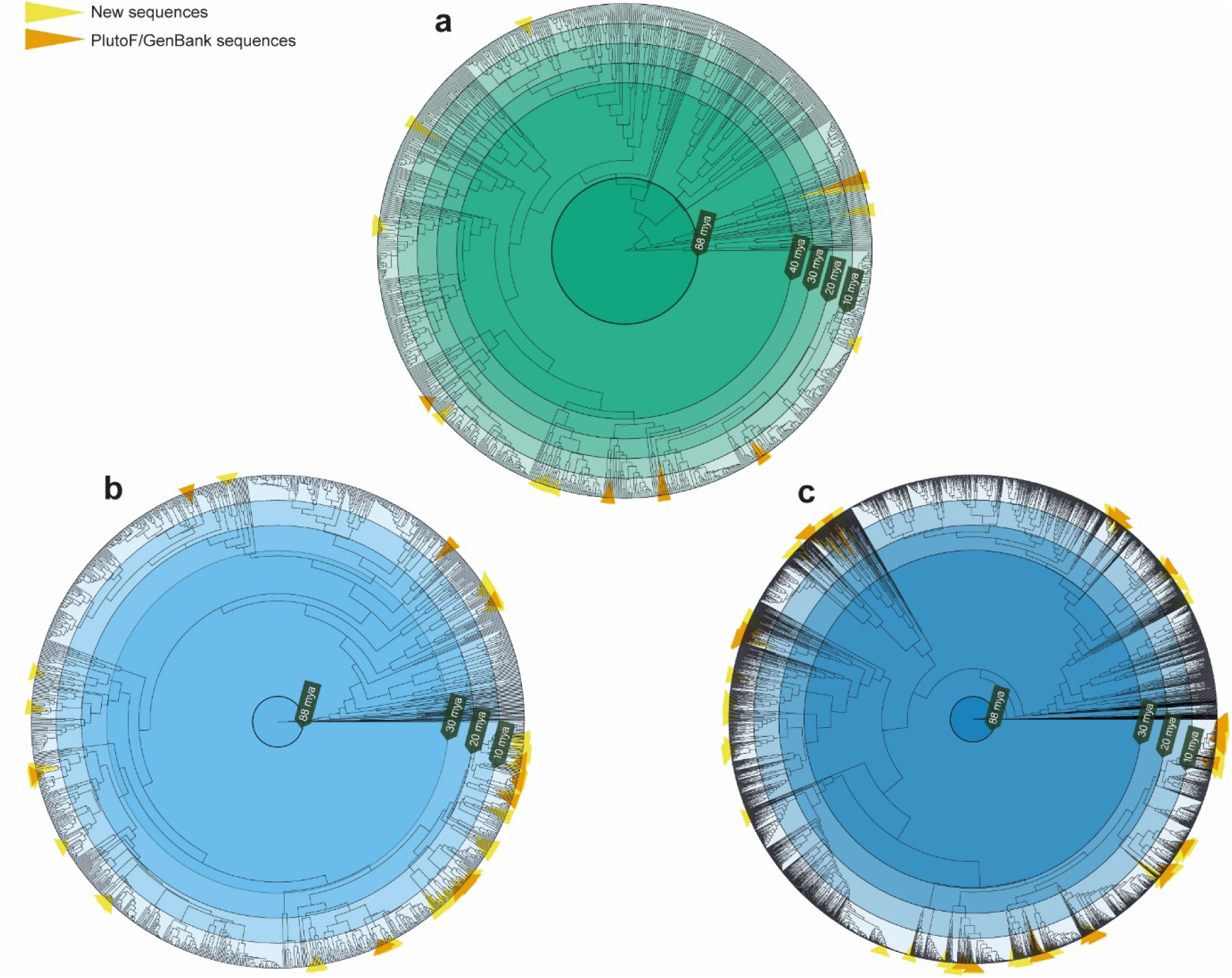
Ultrametric phylogenetic trees for Amanitaceae (a, rooted using 17 sequences from the Amanita sect. Lepidella clade as outgroups), Boletaceae (b, rooted using Chalciporus piperatus as outgroup) and Russulaceae (c, rooted using Gloeopeniophorella convolvens as outgroup). Concentric circles are used to represent different ages, and terminal branches leading to the sequences from Madagascar are highlighted

#### Boletaceae

From the sequences used in the Boletaceae phylogenetic reconstruction, 17 were generated in this study, while 94 sequences were obtained from public repositories. The phylogenetic reconstruction of the Boletaceae family (Fig. 4B) shows that all species from Madagascar diverged from their closest relatives less than 20 mya.

#### Russulaceae

The Russulaceae phylogenetic reconstruction (Fig. 4C) includes 76 sequences generated in this study, and 207 sequences obtained from public repositories. The phylogenetic tree shows that all species from Madagascar diverged from their closest relatives less than 20 mya.

## Discussion

Our study provides the first baseline for characterising the known fungal diversity of Madagascar using ITS DNA sequence data. We combined sequences obtained in opportunistic field surveys with all publicly available sequence data for the fungal DNA barcode region to provide a comprehensive analysis of the known fungal diversity in the island. This approach recovered a higher number of species (620 OTUs) than previously reported (500 OTUs) (Ralaiveloarisoa *et al*. 2016). Given that 10% of the OTUs we recovered were previously unknown, the number of fungal species in Madagascar is likely to increase if new surveys are carried out on the island. In addition to macrofungi, many other important fungal guilds like lichens and endophytes are severely understudied in Madagascar, and identifying them has a high probability of generating many new discoveries.

Our results show that the opportunistic surveys carried out for this study contributed substantially to the known fungal diversity from Madagascar. While it is important to acknowledge that timing and methodology are two important factors that are often overlooked in fungal biodiversity surveys (Halme and Kotiaho 2012), our results clearly show that opportunistic surveys are still effective sources of novel biodiversity in poorly studied areas, such as Madagascar. This highlights the sheer lack of documentation of fungal diversity in Madagascar, as it is the case globally, and reinforces that there is value in small studies and opportunistic surveys in the absence of well-funded and long-term survey projects.

New species are regularly encountered when fungi are surveyed anywhere in the world, whether in well-surveyed places (e.g., Ainsworth *et al*. 2013), through the commercial food trade (Dentinger and Suz 2014, Cutler *et al*. 2020), or poorly surveyed regions (e.g, Castellano *et al*. 2016). Opportunistic surveys continue to be an important source of undescribed species, contributing to the world’s fungal diversity knowledge. Of the 75 newly reported OTUs for Madagascar that we collected in our surveys, 61 did not cluster with any previously published sequence, therefore these are potentially new species to science. A combined taxonomic approach using voucher-based morphology, phylogenetics and ecology to ascertain the novelty and evolutionary position of these OTUs is the gold standard of biodiversity documentation, but will require a substantial commitment to sustained fieldwork with requisite funding.

In our study we identified mainly fungi belonging to Basidiomycota, with the most abundant families being Russulaceae, Thelephoraceae, Boletaceae, Cortinariaceae and Amanitaceae, which are regularly found to be the most abundant ectomycorrhizal clades in tropical forests (Ramanankierana *et al*. 2007; Henkel *et al*. 2012; McGuire *et al*. 2013; Corrales *et al*. 2018).

Most OTUs found in Madagascar could not be identified at species level, meaning that authoritative, unequivocally named ITS sequences for the species they belong to are not available in the online repositories. Sequencing the ITS region of the fungarium specimens described by French mycologists in the late 19^th^, early 20^th^ centuries (Vizzini *et al*. 2019), with the help of high throughput DNA sequencing (Dentinger *et al*., 2016; Andrew *et al*. 2018), would be a step forward in helping naming some of the species we are currently finding in Madagascar, but that we are unable to properly assign a species name.

In our exploration of the EcM diversity of Madagascar, most of the species belong to the /russula-lactarius, /boletus, /tomentella-telephora, /cortinarius and /amanita EcM lineages; these results are in concordance with previously published results of EcM studies from Madagascar (Tedersoo *et al*. 2011, Henry *et al*. 2015). Of all the EcM OTUs recovered, 7% of them came from our opportunistic surveys, mainly belonging to the /russula-lactarius, /amanita, /boletus, /cantharellus, /cortinarius, /sebacina and /elaphomyces EcM lineages.

Interestingly, we found that most of the new diversity revealed in this study came from EcM root-only sequences (60% of all OTUs), with a lower proportion of them coming from sporocarp-only sequences (36%) and with both EcM root and sporocarp sequences (4%). Biodiversity surveys aiming to document unknown species tend to focus their efforts on above-ground sampling of sporocarps (Henkel *et al*. 2012), as one or more sporocarps are needed for a new species to be officially described and published (Aime *et al*. 2021). However, our results show that a significant part of the unknown fungal diversity of Madagascar can be found belowground; therefore, studies aiming to study the overall fungal diversity in tropical forests should sample both above and below the soil surface.

We also found interesting trends regarding the global distribution of the species recovered for Madagascar. Our results suggest that there is a high level of fungal endemism in the island, with 65% of the total number of OTUs only referenced for Madagascar. When exploring the EcM species recovered, the proportion of endemism increases to 81% of the total EcM fungal OTUs. These results agree with previous studies, where a high percentage of fungal morphotypes found in forest from Madagascar could not be assigned to known species, thus potentially being new endemisms (Ralaiveloarisoa *et al*. 2016). Endemism is a consistent result of sequence-based studies of fungal communities at continental and global scales, usually due to climate and environmental variation but also because of dispersal limitations (Peay *et al*. 2016); all of these factors are present in Madagascar. Furthermore, the high levels of endemism found in many other organisms, such as terrestrial plants (depending on the family, ranging from 38 to 100% endemic species, with a mean of 83%), and animal groups such as amphibians (99% endemic), reptiles (92% endemic) and terrestrial invertebrates (86%) (Goodman and Benstead 2005), leads us to believe that it might be the same for fungi.

Our phylogenetic reconstruction shows that all species belonging to the families Amanitaceae, Boletaceae and Russulaceae are much younger than the separation of the India-Madagascar land mass, about 88 mya. All sequences from Amanitaceae diverged less than 34 mya, while for Boletaceae and Russulaceae the divergence occurred less than 20 mya. Our estimates would have to be substantially off for vicariance to be invoked as a primary mechanism for the arrival of EcM fungi in Madagascar.

Both the newly generated sequences and the sequences retrieved from the public databases seem to be phylogenetically overdispersed, with no obvious patterns of clustering between species from Madagascar forming on the phylogenies of the Amanitaceae and Russulaceae. This could be suggesting multiple dispersal events to the island, but this hypothesis needs to be further investigated. Some Boletaceae species from Madagascar are, however, underdispersed over the phylogenetic tree, with clustering on the most derived branches of the tree, in a clade that is still much younger than the separation of the India-Madagascar land mass (88 mya). This suggests a possible role of local adaptation in the diversification of some Boletaceae in Madagascar. However, these interpretations are preliminary and require a larger sampling to provide robust patterns for reliable inference.

Overall, these results do not support our first hypothesis, based on Ducousso *et al*. (2004): if the EcM fungal partners would have arrived in Madagascar before the separation of the India-Madagascar land mass, we would expect to find at least some early-diverging (close to the 88 mya mark) species from Madagascar in the phylogenetic reconstruction of these three EcM families. This was not the case, as we found that all Amanitaceae, Boletaceae and Russulaceae species from Madagascar diverged much later. However, one alternative, albeit highly unparsimonious, interpretation is that while the lineages may have existed prior to separation of the India-Madagascar landmass from continental Africa, they have experience little local diversification but dispersed widely, obfuscating phylogenetic patterns of biogeographic origin. While possible, this scenario would require many unlikely events that at this time seem too improbable to accept given the simpler explanation of a dispersal-driven mechanism.

Moreover, the African and South American Dipterocarpaceae (genera *Monotes* and *Pseudomonotes*, respectively) are now also known to be EcM, which, according to the phylogeny in Ducousso *et al*. (2004),would push the origin of the EcM habit even further back, before the fragmentation of the supercontinent Gondwana between 165-155 mya (Yoder and Nowak 2006). This contrasts with the proposed crown ages for the Malvales, the order that contains both Sarcolaenaceae and Dipterocarpaceae, which range from 63-67 mya to 102.4 mya (Wikström *et al*. 2001, Wang *et al*. 2009, Tank *et al*. 2015, Magallón *et al*. 2015, Hernández-Gutiérrez and Magallón 2019). However, the phylogenetic relationships between Sarcolaenaceae and Dipterocarpaceae are still uncertain (APG 2016), and conclusions should be drawn from their phylogenies with caution.

It can be argued that the relative dating approach used in this study, based on previously published divergence times to date the ultrametric trees (He *et al*. 2019), is fraught with uncertainty. However, secondary calibrations using published divergence times has been used before (e.g., Dentinger *et al*. 2010, Nelsen *et al*. 2020, Zamora and Ekman 2020) in cases where the number of genes or the taxonomic depth of the phylogenies (or both) did not allow for fossil-based dating to be used, which is the case in this study. Notably, our time estimates would have to be off by more than 2-fold (more than 4-fold, in the case of the Boletaceae and Russulaceae), for the EcM families studied to have arrived before the separation of the India-Madagascar landmass; that would push the origin of the EcM families studied to before the origin of the class Agaricomycetes, 298 mya according to He *et al*.’s estimate (2019). Therefore, our estimated dates and corroborating evidence from the estimated dates of the host plants support the arrival of the EcM fungal families Amanitaceae, Boletaceae and Russulaceae in Madagascar well after its separation from mainland Africa.

Given that vicariance is unlikely to be the main mechanism of arrival of the EcM symbiosis to Madagascar, long distance dispersal seems to be the most plausible alternative, and has been proposed for other late-diverging organisms (Renner *et al*. 2001, Nobre *et al*. 2010, Kainulainen *et al*. 2017). Long distance dispersal of EcM fungi is highly variable and across species (Peay *et al*. 2007, 2010, 2012; Geml *et al*. 2012), Wilson *et al*. (2012) described the ancestral distribution of the Sclerodermatineae as North America-Asia-Southeast Asia; they speculate that taxa extant outside of this range (which includes Madagascar) could be attributed to long-distance dispersal capabilities combined with low host specificity. An ancestral area reconstruction of Amanitaceae, Russulaceae and Boletaceae would help decipher the multiple possibilities for the origin of their EcM habit. However, an equal sampling across regions and lineages would be necessary for this approach, which will require major investment since documentation of these diverse families remains highly incomplete. Moreover, using a multi-gene approach would allow for a deeper resolution of the phylogeny (Frøslev et al. 2005, Tremble et al. 2019), allowing for fossil-calibrated dating of the trees that may result in more accurate dates (Zhao et al. 2017, Varga et al. 2019).

Studies on the phylogenies of some of the Malagasy EcM plants are also consistent with long distance dispersal of the hosts. Zanne *et al*. (2014) found that the genus *Uapaca*, present in mainland Africa and Madagascar, diverged from its closest relatives only 14 mya, while Janssens *et al*. (2020) found a similar pattern in the EcM *Intsia bijuga*, diverging from *Afzelia* (Fabaceae) 14 mya. In both cases, the potentially EcM ancestors of these trees would have arrived in Madagascar through long distance dispersal from mainland Africa and the east, either Asia or Oceania (Buyck 2002). While long distance dispersal is not currently known to be exhibited in the zoochorious *Uapaca* (Chirwa and Akinnifesi 2008; Petre et al. 2015), *I. bijuga* is known to use oceanic currents to disperse its seeds (Lewis *et al*. 2005). This may explain the two main ranges of this species in South-East Asia and Oceania, and southern Africa and Madagascar. More ancient long distance dispersal events across the Indian Ocean have also been described by Li *et al*. (2009) where they describe one or possibly two dispersal events in the past 10 million years of *Bridelia* (Phyllanthaceae), an arbuscular mycorrhizal genus with several representatives present in Madagascar.

Our apparently contrasting results of a high level of endemism but EcM speciation through long-distance dispersal suggest that these events are episodic followed by long periods of isolation. This pattern has already been reported for saprobic fungi in the Southern Hemisphere (Moncalvo and Buchanan 2008) and for *Tuber* spp. worldwide (Bonito *et al*., 2010); we hypothesise that the same pattern can be found in the EcM families Amanitaceae, Boletaceae and Russulaceae of Madagascar. Moreover, our Amanitaceae and Russulaceae phylogenies seem to show a pattern of phylogenetic overdispersion of the species from Madagascar, consistent with multiple independent colonisation events, while many Boletaceae species from Madagascar seem to cluster in the furthest branches of their phylogeny, suggesting more limited dispersal events followed by in situ speciation. This difference between the two families also suggests that dispersal events were not synchronous as could happen with, e.g., a major weather event carrying many propagules together. Further research to systematically document EcM fungi and their hosts is needed to test these hypotheses, and to understand the mechanisms of dispersal and the number of times that dispersal events happened across EcM species in Madagascar.

## Conclusion

In this study we provide a first look into the fungal diversity of Madagascar while also highlighting the importance of opportunistic surveys in poorly studied ecosystems. We reconstructed the phylogenies of the EcM families Amanitaceae, Boletaceae and Russulaceae, and found evidence of long-distance dispersal as the most likely mechanism for the origin of EcM fungi in Madagascar.

According to the CBD (2014), more than 80% of the area covered by the original forests in Madagascars has been lost due to destructive human practices such as the clearing of natural habitats and overexploitation of natural resources and this process is still ongoing. Due to the high levels of endemism in the country, the loss of a hectare of forest has a larger impact on its biodiversity than anywhere else in the world. With fungi having been largely overlooked in biodiversity assessments, this could mean that we are losing hundreds of fungal species that have never been described before, and that hold vital information in their DNA about the evolutionary history of the island’s Funga. This is particularly true for EcM species, both plants and fungi, that require a symbiotic relationship between each other to survive. Moreover, EcM fungi are key components of terrestrial ecosystems that play vital roles in plant communities and ecosystem services (Tedersoo *et al*. 2020), therefore decreasing diversity and species loss might result in catastrophic consequences.

Despite the many recent taxonomic studies aimed at identifying new species from Madagascar (Aptroot 2016; Shay *et al*. 2017; Ralaiveloarisoa *et al*. 2020, 2021), our results suggest that a large proportion of its Funga has yet to be described. More thorough fungal surveys are needed in Madagascar, both exploring the existing collections and conducting more frequent above- and below-ground sampling in the field; to achieve this, sustained funding and training of Malagasy mycologists is of paramount importance; this would also help lessen the “helicopter research” (Haelewaters *et al*. 2021) in the island.

## Acknowledgments

This research project was supported by the Queen Mary University of London and Royal Botanic Gardens Kew; When conducting this study, MRF was the recipient of a Fundación Barrié Postgraduate Scholarship Award and a FPU fellowship (FPU20/02280) from the Spanish Ministry of Education; the expeditions to Madagascar were funded by the Kew-Rio Tinto Partnership Fund, Kew’s HLAA Fieldwork Fund and QMUL-Kew’s MSc in Plant and Fungal Taxonomy, Diversity and Conservation. Many thanks to the Kew Madagascar Conservation Centre staff, Alexander Bradshaw, Alexander Byrne, Rowena Hill, Dr. Begoña Aguirre-Hudson, Dr. Sven Buerki, Dr. Felix Forest, Dr. Ricardo Arraiano-Castilho and Dr. Paloma Morán for their help during the sampling, analysing and writing stages of this manuscript.

## Statements & Declarations

### Author Contributions

Sampling was carried out by Laura M. Suz, Bryn T.M. Dentinger, Paul Cannon, Franck Rakotonasolo and Shannon M. Skarha. Sample preparation and DNA extraction and purification was carried out by Laura M. Suz, Bryn T.M. Dentinger and Shannon M. Skarha. Bioinformatic and phylogenetic analyses were carried out by Mauro Rivas-Ferreiro. The first draft of the manuscript was written by Mauro Rivas-Ferreiro, Laura M. Suz and Bryn T.M. Dentinger, and all authors commented on previous versions of the manuscript. All authors read and approved the final manuscript.

### Data availability

The new sequences generated in this study are available in GenBank under the Accession Numbers: (insert here). All other datasets generated during and/or analysed during the current study are available from the corresponding author on reasonable request.

## Supplementary Information

ESM_1.xlsx: **Table S1**. List of GenBank accession numbers for all sequences generated in this study

ESM_2.xlsx. **Table S2**. List of OTUs found in this study from Madagascar, including the *dada2* identification, the SH (if available), the known distribution and the EcM lineage.

